# A pancrustacean brain and its visual systems define a Cambrian “great appendage” arthropod

**DOI:** 10.1101/2025.01.24.634743

**Authors:** Nicholas J. Strausfeld, Xianguang Hou, Frank Hirth

## Abstract

Early Cambrian fossils from the Chengjiang biota demonstrate that over half a billion years ago early stem euarthropods existed coevally with representatives of already recognizable crown groups. Prominent stem taxa were *Fuxianhuia protensa* and *Alalcomenaeus* whose cerebral and ganglionic traits identify them as, respectively, stem mandibulates and stem chelicerates. Here we report features of the visual systems and brain of the enigmatic lower Cambrian stem euarthropod *Jianfengia multisegmentalis*, which despite the absence of trunk tagmatization reveals neural traits that justify its inclusion within stem Pancrustacea also known as Tetraconata. The sutured eyestalks of *Jianfengia,* typifying crown Malacostraca, terminate as compound eyes populated by ommatidia that reveal tetradic structures indicative of cone-building Semper cells. Preserved neuropils and tracts in the eyestalks of *Jianfengia* resolve nested optic neuropils as occur in eucrustaceans. The nauplius-like eyes of *Jianfengia* and their associated nerves supply a discrete forebrain region, as they do in pancrustaceans. These attributes distinguish Pancrustacea from stem Chelicerata and extant Myriapoda as does the organization of the jianfengiid deutocerebrum and tritocerebrum. In the absence of external traits that differentiate identities amongst stem euarthropods, neuroanatomical traits provide a powerful tool for discerning characteristic distinctions and correspondences for elucidating euarthropod relationships. Here, we demonstrate that cerebral arrangements in *Jianfengia* correspond to those of pancrustaceans existing today.

## Introduction

Fossils of stem euarthropods of the lower Cambrian exhibit neuromorphological characters that typify extant crown Euarthropoda (1, 2): the organization of the optic tracts and neuropils subtending the compound eyes of *Fuxianhuia protensa* corresponds to that of visual centers defining mandibulates; organization of the fuxianhuiid deutocerebrum with its antenniform appendages (3) corresponds to that of mandibulate olfactory lobes and antennules; brain and visual systems of the megacheiran *Alalcomenaeus* reveal a cerebral organization corresponding to that of larval *Limulus* (Merostomata) (4). Such distinctions relate directly to variations of neural organization within each of the three domains of the cerebrum that in extant euarthropods are genetically determined by the combinatorial activity of conserved homeobox transcription factors, thereby indicating their ancient origin (5). Specific phenotypic distinctions of the internal organization of each cerebral domain further discriminate the divergent euarthropod clades Chelicerata, Pancrustacea, and Myriapoda. Unrecognized until now are fossilized species that possessed cerebral traits corresponding to those that define today’s most species-rich group, the Pancrustacea (6).

Here we describe observations of three specimens of the genus *Jianfengia multisegmentalis* (Hou 1987) (7) retrieved from the Cambrian (Series 2, Stage 3) Eoredlichia–Wutingaspis trilobite biozone, Yu’anshan Member, Chiungchussu Formation. Unlike previous euarthropod fossils that have revealed neural traces indicative of brains millimeters wide (3, 4), specimens of *J. multisegmentalis* are uniformly minute. A carapace barely 1 mm wide offers evidence of putative brains that are an order of magnitude smaller than, for example, that of *Fuxianhuia*. Neural traces resolved with high magnification require optical and digital adjustments to demarcate shapes and trajectories. We report two specimens of *J. multisegmentalis* whose shapes and trajectories of fossilized neural traces align with brains of extant eucrustaceans: rostral nauplius eyes and brain areas receiving their axons; paired eyestalks capped by compound retinas whose ommatidia possessing tetrad arrangements of cone cells; and nested optic neuropils connected to an asegmental cerebrum leading to the first true segmental ganglia. Together these findings identify *Jianfengia* as a stem euarthropod typifying pancrustaceans and distinguish it from both *Fuxianhuia* and *Alalcomenaeu*s.

## Results

### Sensory attributes and diagnostic traits

As shown in specimen YKLP11117 (Fig. 1A), a jianfengiid’s trunk can reach 2-2.5 cm in length with another 5 mm taken up by the rostral carapace barely1mm in width. The carapace covers the first four trunk segments and the cerebrum except for certain elements of the rostral sensory systems. These are the laterally extending eyestalks of the protocerebrum and the most rostral cephalic cuticle whose paired anterior projections flank a medial anterior sclerite, which is separately attached to and articulates with the exoskeleton (Fig. 1B-D). Despite being flattened, the anterior sclerite suggests a volume that is capped by a system of lenses. Their detailed organization suggests structural homology to crustacean nauplius eyes (8, 9) and insect ocelli (10, 11) (Fig. 1C, D; *SI Appendix* Fig. S1) comprising three lenses overlying a palisade of photoreceptors.

**Fig. 1.**
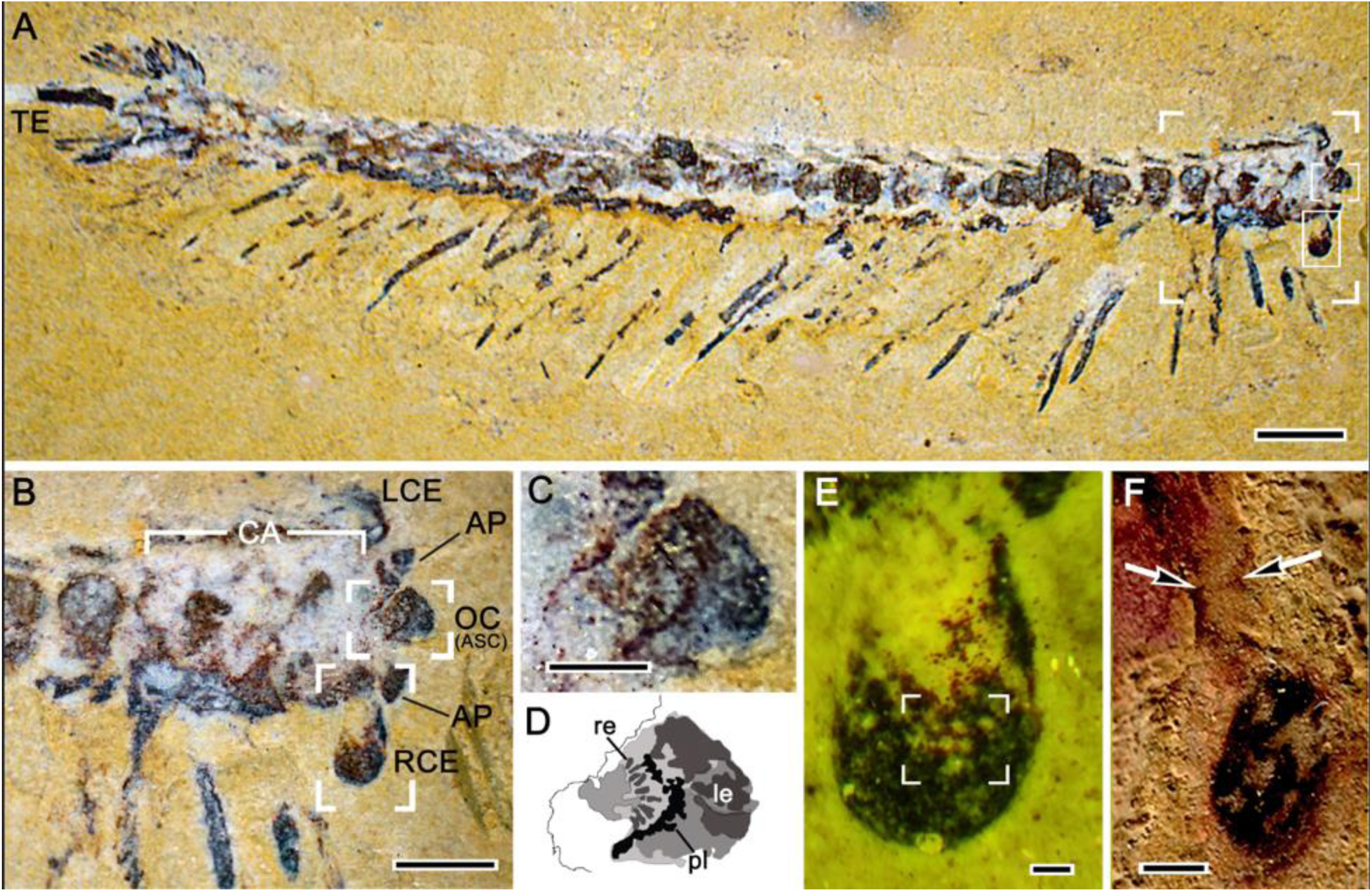
The ocellar and compound eye visual systems of *Jiangfengia*. (**A**) Top-down view of YKLP11117. Its trunk, composed of 27 isomorphic segments, terminates in an elongate telson (TE). The framed area to the right, which includes the carapace and cerebral elements, is enlarged in panel b. Within the frame, the upper box indicates the ocellar nature of the anterior sclerite enlarged in panels B-D. (**B**) The four rostral-most segments and the asegmental cephalon are covered by the carapace (CA) extending forward from the 4^th^ trunk segment. Eyestalks situated behind the paired anterior projections (AP) emerge laterally from the front of the carapace (right eye RCE, fully exposed) and terminate as compound eyes (LCE, RCE). The midline ‘anterior sclerite’ (ASC) carries the ocelli (OC; see *SI Appendix,* Fig. S1). (**C, D**) The ocelli: a palisade of rod-like elements (re) beneath a layer of darker pigment (pl) indicates the likely ocellar photoreceptor layer. **(E)** Right compound eye, photographed under combined UV and white light, showing some facets with missing lenses (box) allowing the resolution of Semper (cone) cells, as shown in Fig. 2. (**F**) A distinct suture (arrowed) of the eyestalk indicates its constituent podomeres. Scale bars: A= 2mm; B =1mm; C, F=0,25mm; E =50 μm.

Superimposition of neural traces from the anterior sclerite of specimen YKLP17299 onto the corresponding exoskeleton of YKLP11117 demonstrates a system of axons from the nauplius eyes/ocelli extending back into the most rostral neuropil of the brain (*SI Appendix,* Fig. S1). As has been shown by developmental genetics of extant pancrustaceans, the combinatorial activity of *Six3, FoxQ2* and *hbn* gene homologs is required for the development of the rostral ocellar photoreceptors whose axons project into the prosocerebrum of extant pancrustaceans (12–14). The corresponding organization of nauplius eyes in *Jianfengia*, including their central axonal projections (Fig. 1A-D; *SI Appendix,* Fig. 1), likewise identify its prosocerebral domain. Conspicuously absent in extant Myriapoda (and in Fuxianhuiida), this nauplius/ocellar visual system is restricted to Pancrustacea (15).

*Jianfengia* possesses compound eyes on paired eyestalks that extend laterally from beneath the carapace (Fig. 1B, E). The eyestalks correspond to those typifying eumalacostracan crustaceans (16) in that each stalk evinces two articles connected by a suture (Fig. 1F). In extant taxa, the combinatorial activity of *Six3*, *Otx* and *Pax6* defines the protocerebral domain of the pancrustacean brain (12, 17) and its nested optic neuropils which receive information from the eye’s ommatidial array (18). Compound eyes on stalks and their associated optic neuropils thus indicate this forebrain domain of *Jianfengia* to be the protocerebrum.

A trait that exclusively defines pancrustaceans is further demonstrated by the compound eyes of *Jianfengia*. The loss of some cuticular lenses in specimen YKLP11117 allows a view into their underlying ommatidial shafts. Illumination of these by combined UV and white light reveals a geometric arrangement of internal elements (Fig. 1E; Fig. 2). To determine if these are taphonomic artifacts or that they represent fossilized cellular elements of the ommatidia, we applied an Adobe Photoshop function that resolves peak intensities in a defined chromatic range (Fig. 2; see Materials and Methods). This reveals the iridescent structures shown in Fig. 1E to be elements of an ommatidium, each group organized as a quartet (Fig. 2, rows D, E, F1-4). The same function applied to areas of the matrix in which the specimen is embedded identified peak intensities whose structure bears no resemblance to the ommatidial quartets (Fig. 2, column 5). In extant pancrustaceans, this tetrad arrangement corresponds to only one cell type amongst those that comprise an ommatidium. These are the Semper cells, crucial in the patterning of the compound eye and the formation of its ommatidia (19). In pancrustaceans four Semper cells develop at the base of an ommatidium, each cell sending out a single extension that contributes to the ommatidium’s secreted crystalline cone, which with the overlying cuticular lens provides the ommatidium’s dioptric apparatus (20). Scutigerid centipedes of extant Myriapoda uniquely possessing compound eyes (21) also have quartets of Semper cells but these are distinct from those of pancrustaceans in that each of the quartet contributes at least two extensions that contribute to the provision of the crystalline cone (22). The fact that in insects and crustaceans each cone comprises four parts each part determined by a single Semper cell prolongation has inspired the term ‘Tetraconata’ as the alternative name for Pancrustacea thereby emphasizing the inclusion of hexapods and eucrustaceans within a single clade (23).

**Fig. 2.**
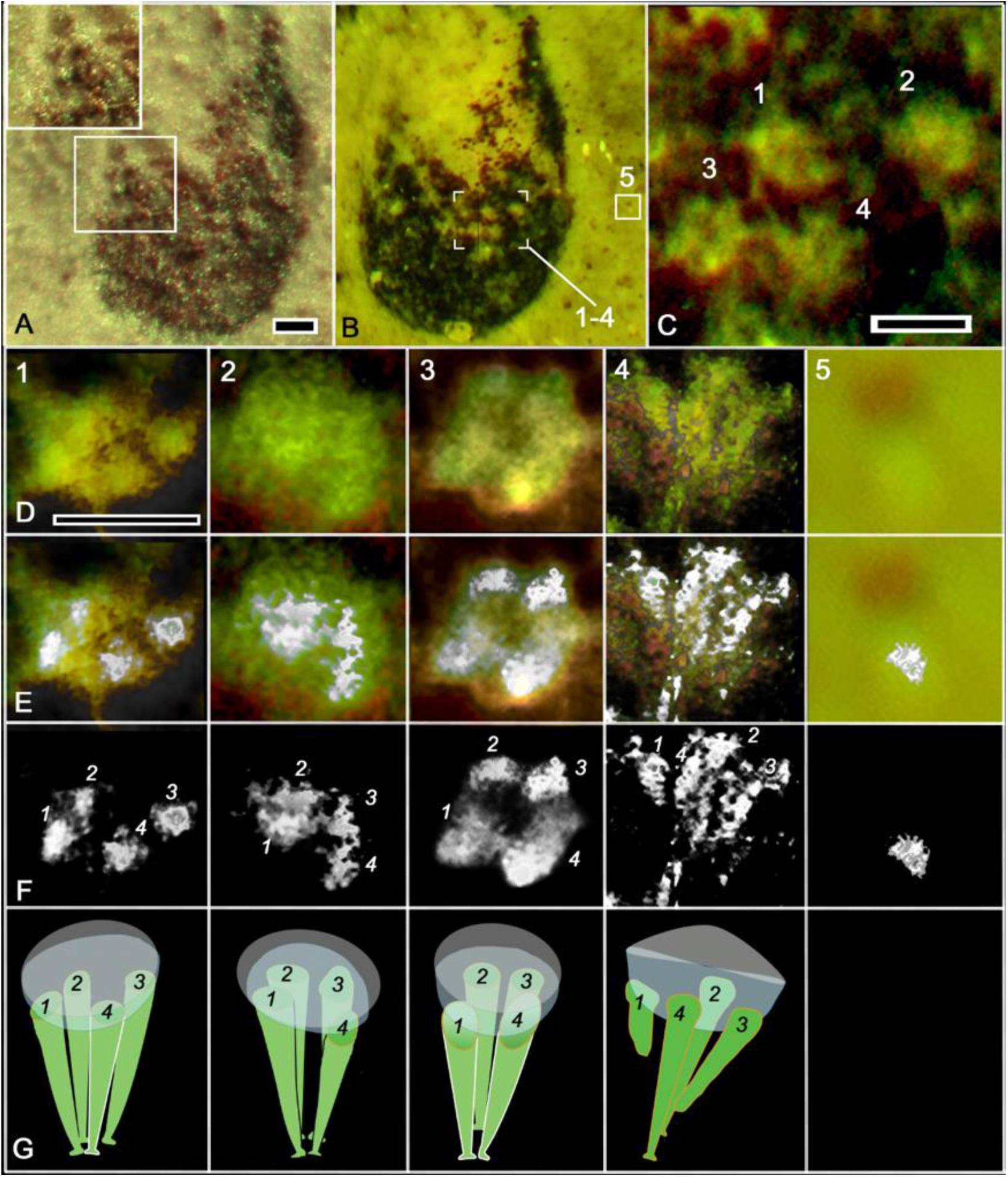
*Jianfengia* compound eye and its tetraconate organization. (**A**) The compound eye of specimen YKLP11117 comprises approximately 120 facets some of whose convex lenses (enlarged upper left) indicates their hexagonal arrangement. (**B, C**). The boxed area in panel B indicates four ommatidia (1-4 enlarged in panel C) whose lenses have been lost, thus revealing structures beneath (1-4 in panel C). Rectangle 5 in panel B indicates one of several bright areas of the matrix sampled to exclude spurious taphonomic-generated patterns. (**D1-4**) Enlarged images within ommatidia 1-4 in panel C. (**E1-4**) Images overlain by their computed maximum intensity levels. (**F1-4**) Maximum intensities alone demonstrate four profiles in each ommatidium. (**G1-4**) Interpretive diagrams of tetrad Semper (cone) cells extending outwards to the crystalline cone (light gray) that would have been overlain by its lens (darker gray). Numbers indicate the orientations of each quartet of Semper at their observed focal plane. (**column 5**) Intensity values at control area 5 in panel B. Scale bars: 50 μm in panel A; 20μm in panel C; 20μm in panel D for rows D-F.

### Optic lobes and brain

We were able to identify cerebral neuropil in the minute specimens YKLP11367 and YKLP17299. In both, reconstruction of brain and nervous system underlying the carapace (Fig. 3A-I) was achieved by processing high-definition images to eliminate asymmetric imbalance and chromatic discontinuities that typify Changjiang fossils as described in the methods section and its supporting in formation (see Materials and Methods, below with *SI Appendix* Figs. S3, S4). The resultant reconstruction (Fig. 4) resolves cerebral neuropils and associated ganglia whose dispositions correspond to traits typifying the brains described for stemward groups (6) that include Cladocera, Anostraca and Decapoda (24–26). As described for the fossilized cerebrum of Leanchoiliidae (27), traces of neural tissue within the jianfengiid stomodeum (Fig. 3B, D, I) align with the location of the appendicular labrum (Fig. 3E) resolved by micro-CT imaging of the underside of specimen CJHMD 00061 (see Acknowledgements). A second specimen of *Jianfengia* (YKLP17299) provides evidence of fossilized axon bundles diverging from ocellar/nauplius eyes to the forebrain where they merge with bilateral neuropils situated extreme rostrally in the prosocerebrum (Fig. 3F; also *SI Appendix,* Fig. S1). The divergent trajectories of these nerve cords are identical to those described for crustacean nauplius eyes and insect ocelli (8–10) which reach discrete neuropils specific to the pancrustacean prosocerebrum (11).

**Fig. 3.**
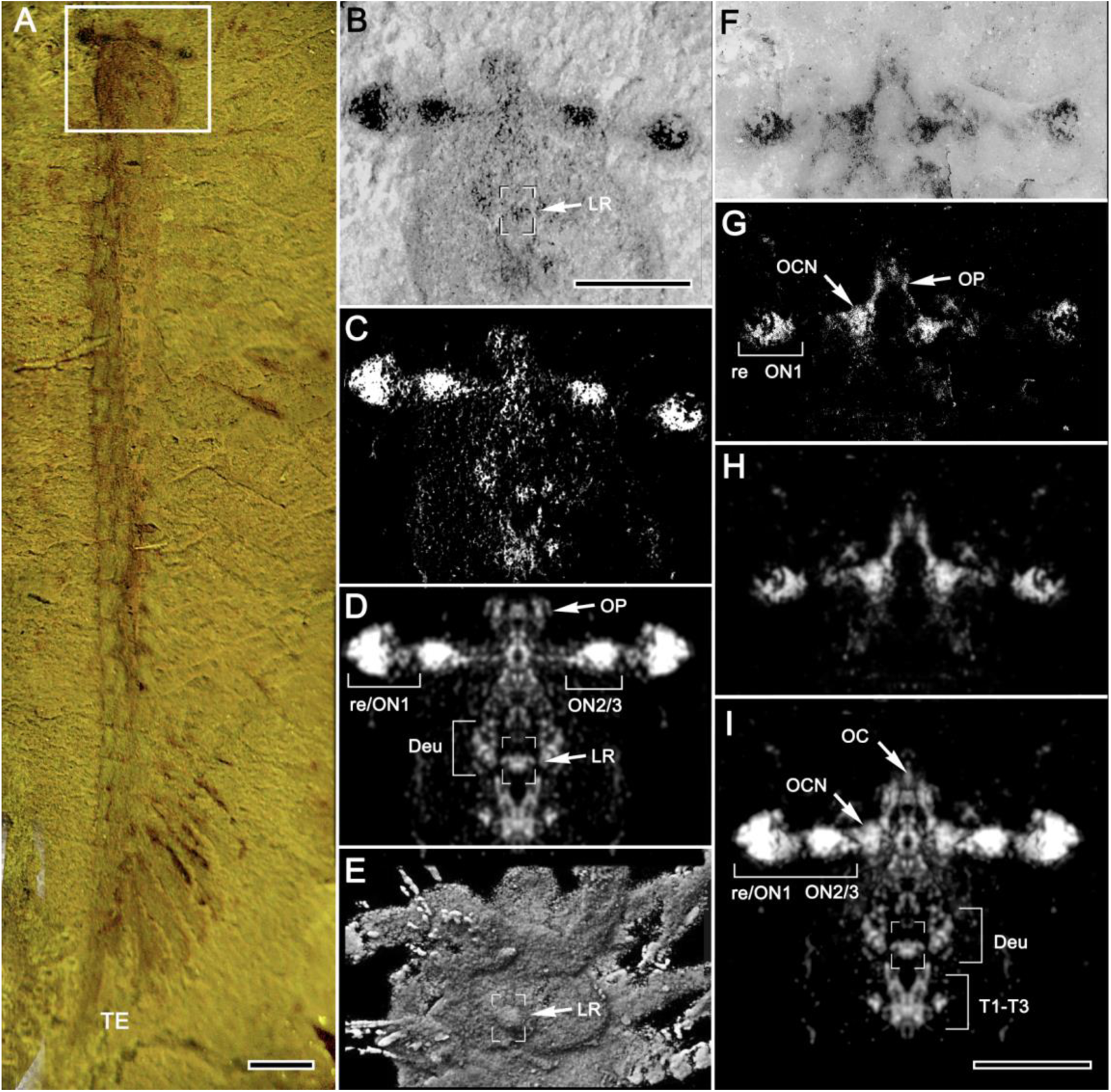
The brain of *Jianfengia*. Reconstruction of fossilized neural traces, neuropils, and connections of specimen YKLP11367 and YKLP17299. (**A**) Boxed area indicates the area providing neural traces processed in panels B-D. (**B**) Gray level image of the cephalic area. Traces of neuropil and tracts are indicated by darkest deposits. The dark profile within the stomodeum indicates paired labral ganglia (box LR), also indicated in panels D and I, and matching the cuticular labrum shown in panel E. **(C)** Raw extracted image of neural traces. **(D)** Bilaterally symmetric rendering of neural traces (see Extended Data Fig. 3 for reconstruction steps). **(E)** Micro CT image of the underside of specimen CJHMD 00062 (see acknowledgements) showing the labrum (boxed) aligned with neural traces likewise indicated in panels B, D and I. **(F)** Anterior of specimen YKLP17299 showing well-defined ocellar tracts from the anterior sclerite supplying the rostralmost forebrain. **(G)** Raw extracted image of neural traces. **(H)** Panel g subjected to Gaussian blur. (**I**) Overlay of panel D and H provides composite image of the jianfengiid brain and its subsequent three segmental ganglia arranged as a contiguous synganglion. Abbreviations: TE, telson; ON1-ON3, volume and profiles indicating 3 contiguous optic neuropils (9) proximal to the terminal retina (re); OP, ocellar pathways; OCN, discrete rostro-lateral neuropils; Deu, deutocerebral domains lateral to the stomodeum; T1-T3, post oral ganglia arranged as a synganglion. Scale bars: A=1mm; B-I=0.5 mm.

**Fig. 4.**
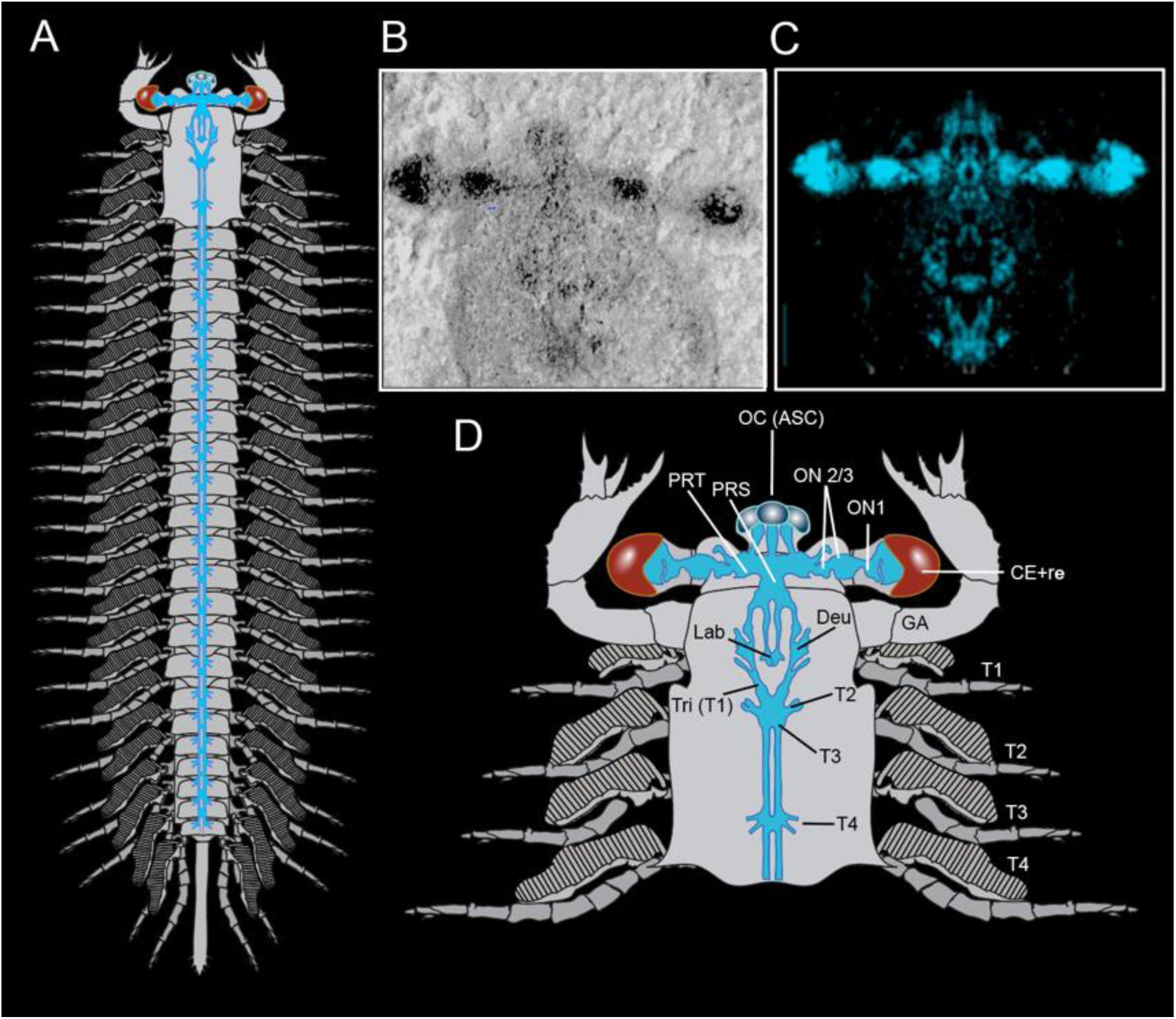
Reconstruction of *Jianfengia multisegmentalis*. (**A**) Total idealized reconstruction based on specimen YKLP11117, YKLP11367 and YKLP17299. Segmental ganglia have not been resolved caudal to T3. Those shown are hypothetical, based on documented examples in other euarthropods (27). Trunk appendages are biramous: pairs beneath the carapace have 5-7 articles, the rest have 10, except the last few (28). (**B**) Fossil neural tracts; (**C**) reconstructed brain and post-oral synganglion. (**C**) Annotated interpretive reconstruction of the cephalic area. Abbreviations: Deu, deutocerebrum; GA, great appendage; Lab, labrum; OC (ASC), ocelli, anterior sclerite; ON, optic neuropil; CE+Re, Compound eye+retina; PRS, prosocerebral domain; PRT, protocerebral domain.

The merged images of YKLP11367 and YKLP17299 (Fig. 3I) provide a composite reconstruction of the jianfengiid brain and its connecting ventral nerve cords. The eyestalks of YKLP11367 (Fig. 3D) reveal the distal retinal layer (re) and its small first optic neuropil (re/ON1) contiguous with a voluminous visual neuropil (ON2) adjacent to a smaller one (ON3). These nested arrangements, which are exclusive to pancrustaceans (18), are connected to the protocerebrum centripetally by a prominent optic tract (Fig. 3D, I). The protocerebrum gives rise to caudally directed connections that extend part way around the esophageal foramen where they then swell into bilaterally situated deutocerebral neuropils (Deu in Fig. 3D, I). The nerve cords extending further caudally from each half of the deutocerebrum converge post-orally to connect with three fused ganglia (T1-T3 in Fig. 3I). Whereas this synganglion is well resolved, only a satellite trace suggests clustered afferent axons dispersing towards the first three pairs of trunk appendages. A complete reconstruction of the fossil with its cerebrum is shown in Fig. 4, including the assumed 1:1 disposition of the trunk’s segmental ganglia.

## Discussion

The present data identify cerebral and visual system traits that categorize *Jianfengia* as a lower stem pancrustacean even though its three post-oral trunk segments show no differentiated appendages except for a shortening of the first three pairs (28). In crown eucrustaceans the specialized first trunk segment (T1, the tritocerebrum) is recognized as the anterior limit of Hox gene expression and the restricted expression of the gene *collier* (29). Although in extant euarthropods T1 becomes integrated into the brain as a result of morphogenetic movements, it is genetically distinct from the forebrain (ce1-ce3) and demarcates the interface between the asegmental cerebrum and the segmental ganglia of the ventral nervous system (*SI Appendix,* table S1). The second trunk segment T2 of *Jianfengia* corresponds to the extant pancrustacean mandibular segment, but in *Jianfengia* morphological traits expected of mandibles are completely absent whereas in crown pancrustaceans, their patterning and differentiation depend on region-specific expression of the transcription factor *cap’n’collar (cnc)*, which is restricted by the anterior Hox gene *Deformed (Dfd)* (30–32). Possibly, the homonomy of trunk appendages in *Jianfengia* indicates the absence of early differential Hox gene activity underlying trunk tagmatization (33). Yet the coalescence of the first three of *Jianfengia*’s segmental ganglia (Fig. 3, 4) speaks against this, as does the reduction of podomeres in the corresponding first three biramous appendages (28). The arrangement in crown pancrustaceans of fully differentiated tritocerebral, mandibular, and maxillary appendages must have emerged at some time in early euarthropod evolution. That in *Jianfengia* the tritocerebral ganglion is grouped together with segments T2 and T3 as a synganglion ((Fig. 4) also suggests that differential Hox gene activity was already in play. This organization of a tritocerebrum contiguous with the mandibular and first maxillary ganglia also distinguishes the jianfengiid nervous system as radically distinct from that of Myriapoda where the intercalary tritocerebral segment in extant groups can be positioned so far forward as to be almost assimilated into the myriapod deutocerebrum (34).

If traits defining its cerebrum distinguish *Jianfengia* as a lower stem pancrustacean, then why aren’t its deutocerebral appendages antenniform? Instead, they are robust pincer-like structures termed “great appendages” (*SI Appendix,* Fig. S2). These have historically been attributed to diverse stem euarthropods grouped into the clade Megacheira (35, 36). Phylogenetic analyses relying on exoskeletal attributes favor Megacheira as paraphyletic rather than representative of stem Chelicerata, which has its own uncertainties (37, 38). Yet when considering their cerebral and segmental traits, at least two natural groups become apparent (*SI Appendix,* table S2). One comprises *Alalcomenaeus* and *Leanchoilia*, both accepted as lower stem Chelicerates (4, 40), which are characterized by fourteen homonomous trunk segments, paired deutocerebral “great appendages” comprising elongate pincers whose chelae provide annulated flagellae, and cerebra supplied by four semi-sessile limulus-type eyes (4, 29). The other megacheiran group includes *Jianfengia* with two other genera, *Pseudoiulia cambriensis* and *Fortiforceps foliosa* (39, 40) the trunks of which comprise twice as many homonomous segments. All of the second group possess compound eyes, but *Jianfengia* is thus far only genus identified with ocelli and eucrustacean-type eyestalks (Fig. 4). The “great appendages” of *Jianfengia* are stubby plier-like chelae (*SI Appendix,* Fig. S2A-C). The plausibility that a lower stem pancrustacean could be equipped with stubby great appendages rather than antennules should not be readily dismissed, however. Developmental genetic analysis demonstrates that interference with the genetic program defining the formation of aranean chelae – extant homologues of great appendages (41) – would lead to a genetic reorganization of the deutocerebral appendage resulting in a switch from the deutocerebral “great appendage” morphology to that of a deutocerebral antennule (42). Both kinds of appendages, chelae and antennules, are uniramous and although stressed as distinct, there are noticeable likenesses between “great appendages” and certain antennules in that both are characterized by “double axes” (43). Even now there are eucrustaceans that develop great appendage-like antennules, as in male harpacticoid copepods, where antennules terminate as two opposing podomeres the function of which is to constrain a juvenile female until she is fertile enough for insemination (44).

Traits of the fossilized brains of the stem euarthropods *Fuxianhuia protensa* and *Alalcomenaeus* distinguish, respectively, their mandibulate and chelicerate identities. As shown in Fig. 5, the brains of *Alalcomenaeus,* and *Jianfengia* align with the brains of extant Panarthropoda based on the conserved relative order of cerebral domains, their associated appendages, and sensory modalities. In extant Euarthropoda, each cerebral domain is defined by the combinatorial activity of conserved homeobox transcription factors, with *Emx, Nk2* and *Exd* defining the ce3 domain and its deutocerebral integration centers, which in Cambrian stem euarthropods equally serve the ‘great appendages’ or their homologous antennules (5). Thus, it is not Hox gene-associated divergences of the segmented trunk that invariably align ancestral and crown euarthropods; rather, it is their asegmental cerebral domains, their unique lineage-specific neuropils and associated sensory traits (Fig. 5) that define the ancestral status of lower Cambrian stem euarthropods. Here we have shown that amongst those hereditary taxa *Jianfengia multisegmentalis* provides the most stemward evidence of the pancrustacean/tetraconate identity.

**Fig. 5.**
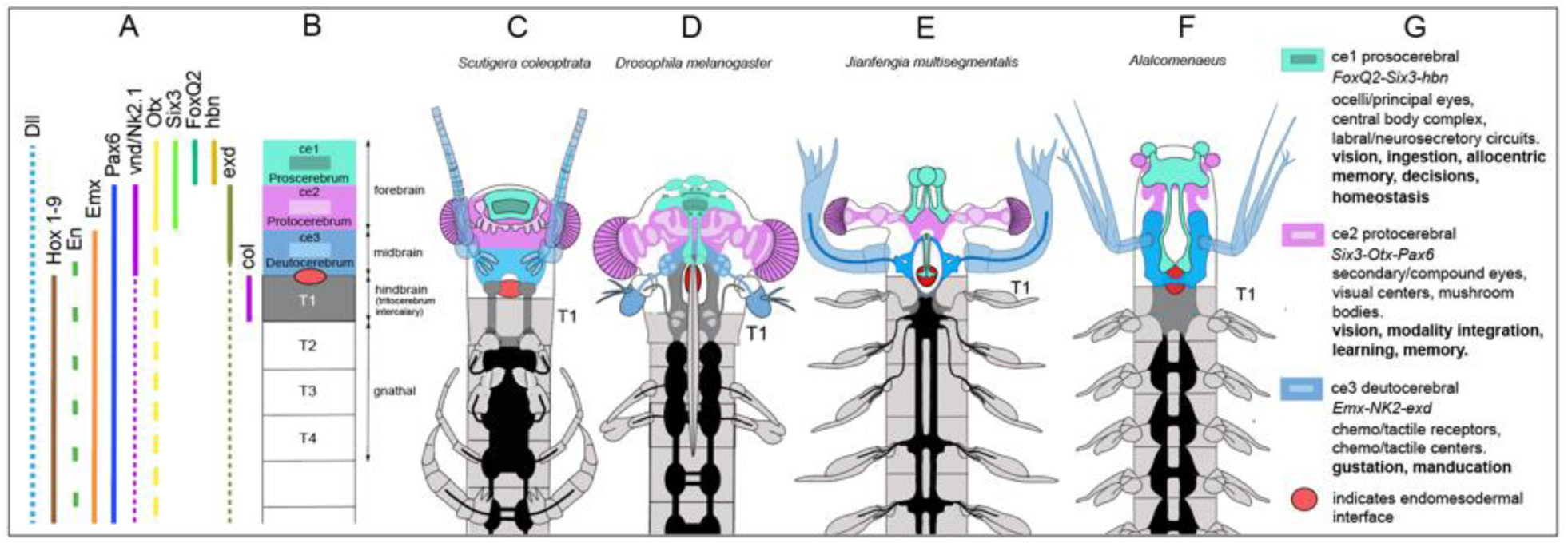
The brains of fossilized Megacheira align with those of extant Mandibulata. (**A, B**) Combinatorial expression of homologous genes in living euarthropods defines three cerebral domains: ce1 (prosocerebrum); ce2 (protocerebrum); ce3 (deutocerebrum) (5). T1 is the first true trunk segment specified by *collier* (29). The interface between the asegmental brain and segmented trunk coincides with the interface between the fore- and midgut, referred to as the endomesodermal interface (EMI, red ovoid). When conserved in fossils, evidence for the EMI provides a non-neural reference point allowing alignments as shown here. Appendages of segments T2-T4 evolved for feeding are referred to as gnathal. (**C**) Schematized rostral nervous system of Chilopoda (Myriapoda). Its ce1 domain is denoted by the central complex. No members of Chilopoda possess a prosocerebral visual system. Two small optic neuropils in the ce2 domain of scutigerid chilopods receive afferents from paired compound eyes. The ce3 domain’s afferent supply is provided by elongated antennules. The intercalary tritocerebrum (segment T1) is pierced by the gut; like in Diplopoda, each of its small hemiganglia are almost engulfed by the deutocerebrum (34). (**D**) In pancrustaceans (here represented by *Drosophila*) the ce1 domain receives ocellar afferents, contains the central complex, and provides the labral “ganglia” via the median bundle (12–14, 45). The ce2 domain is denoted by faceted tetraconate eyes, large nested visual neuropils, and mushroom bodies. The ce3 domain and T1 neuropils receive diverging chemo- and mechanosensory inputs from the ce3 antennules. In Hexapoda, T1 is intercalary lacking second antennae. (**E**) The ce1 domain of *Jianfengia* is defined by its nauplius-like/ocellar visual system and rostral neuropils. The c2 domain is denoted by its tetraconate compound eyes and nested optic neuropils. The ce3 domain flanks the anterior part of the stomodeum as swollen bilateral neuropils serving the “great appendages.” Descending nerve cords from ce3 meet at a synganglion comprising the first three segmental ganglia T1-T3; connections from these and from the T4 ganglion are conjectural. (**F**) *Alalcomenaeus* is recognized as an upper stem chelicerate (4). Its paired medial eyes are prosocerebral homologues of pancrustacean nauplius eyes. Lateral single-lens eyes supply neuropils of the protocerebrum (ce2). Labral ‘neuropil’ originates from the prosocerebrum (ce1). (**G**) In extant Euarthropoda, each cerebral domain is defined by the combinatorial activity of conserved homeobox transcription factors that also specifies domain-specific neuropils and sensory-functional modalities (5, 45). This relative order, including associated appendages, is conserved in fossilized brains.

## MATERIALS AND METHODS

### Provenance

Descriptions herein refer to specimens of *Jianfengia multisegmentalis* retrieved from the Cambrian (Series 2, Stage 3) Eoredlichia-Wutingaspis trilobite biozone, Yu’anshan Member, Chiungchussu Formation, Haikou. The specimens YKLP11117, YKLP11367, YKLP17299 and NIGPAS 100123b are curated at the Yunnan Key Laboratory for Palaeobiology (YKLP), Institute of Palaeontology, Yunnan University, Yunnan, Kunming, China.

### Photomicroscopy

For light microscopy, digital images of the fossils were taken using a Nikon D3X attached to a Leica M205C photomicroscope (Leica Microsystems; Wetzlar, Germany). Images were transferred to Adobe Photoshop CS5 (Adobe Systems; San Jose, CA) and processed using the Photoshop camera raw filter plug-in to adjust sharpness, luminance, texture, and clarity. For ultraviolet illumination, fossils were photographed using a Leica MZ10 F stereomicroscope with appropriate filter blocks to evoke intense green fluorescence. Flexible fiberglass light guides were used to combine UV fluorescence and white-light illumination.

### Identifying tetraconate Semper cell arrangements

Enlargements of the compound eye of YKLP11117 shows its composition of about 120 facets. Patches of these at the eye margin and about 2/3rds across resolve their arranged hexagonal organization. (Fig. 2A, B). Four ommatidia (numbered 1-4 in Fig. 2B) show a loss of their capping lenses thereby revealing structures within the ommatidial shaft. We used ultraviolet illumination to reveal clusters of bright green images within four ommatidia (numbered 1-4 in panel C). These configurations were then resolved at high magnification and are shown in columns 1-4 with steps in their analysis and reconstruction in rows D, overlain in row E by their computed maximum intensities shown in row F). Four profiles (numbered 1-4) reside in each sampled ommatidium. These allow their interpretation in row G as tetradic arrangements of Semper (cone) cells extending outwards to the crystalline cone (light gray) that would have been overlain by its lens (darker gray). The reconstructions of Semper cell tetrads in row G are extrapolations centripetally from the dispositions of their optical sections in row F (represented as 1-4 in row G) to their assumed origin at the inner margin of the retina in row in row G. To determine whether these tetrads might be chance configurations reflecting inconsistences of the embedding matrix the same procedure used to image within ommatidia was used for sampling outside the fossil as in the area denoted as “5” in panel B with intensities and patterns shown in columns Fig. 2B.

### Image processing of detected neural traces

Selective removal of specific tints resolves cryptic traces of fossilized soft tissue but also enhances the contrast of particulate surface features (*SI Appendix* Fig. S3A-B). These imbalances across the specimen are rectified by converting the image to its greyscale mode, merging grey level densities responsible for imbalance and adjusting this to provide approximate uniformity across the specimen (*SI Appendix* Fig. S3C-F). Adjustments of exposure and offset provide the final image for analysis and reconstruction, as shown in Fig. 3B. Adobe Photoshop functions eliminating intensity levels beneath a defined threshold (here 87% black in the standard CMYK scale) provided features that were first reversed as white profiles on a black background (Fig. 3C, G) and then subjected to a Gaussian blur function (R=10px) to provide resolution of neural residues free of background noise (Fig. 3D, H). To establish a bilaterally symmetric reconstruction of the brain and to account for the asymmetry of neural traces around the midline, we used the left side as the image base and flipped the traces from the right side (excluding the eyestalk) over onto it, merging the two halves, using the stomodeal midline for alignment (*SI Appendix* Fig. S4D-G). Two identical brain halves were then digitally generated and joined mirrored symmetrically as shown in (Fig. 3D, H) followed by superimposition of the image of the second specimen YKLP17299 (Fig. 3F-G) with that of YKLP11367 (Fig 3H).

## ACKNOWLEDGEMENTS

We thank Ms. Ziying Wu for photography assistance, and Dr. Yu Liu for computed tomography images of specimen CJHMD 00062 showing the *Jianfengia* labrum/hypostome complex. Our special thanks go to Dr. Camilla Strausfeld for her advice and critical editing of the manuscript’s penultimate version. Dr. Wulfila Gronenberg gave pivotal advice regarding appropriate deployment of Adobe Photoshop functions. This work was supported by the National Science Foundation under grant 1754798 awarded to N.J.S. F.H. acknowledges support from the UK Biotechnology and Biological Sciences Research Council (BB/N001230/1).

## AUTHOR CONTRIBUTIONS

N.J.S. and F.H. originated the project. X.H. discovered the species, provided specimens and with N.J.S. discussed the material. N.J.S and F.H. wrote the manuscript. F.H. ascribed published gene expression data to described panarthropods.

## SUPPORTING INFORMATION

### Tetraconate neural identity defines a Cambrian great appendage euarthropod

Figs. S1 to S4

Tables S1-S2

**Fig. S1.**
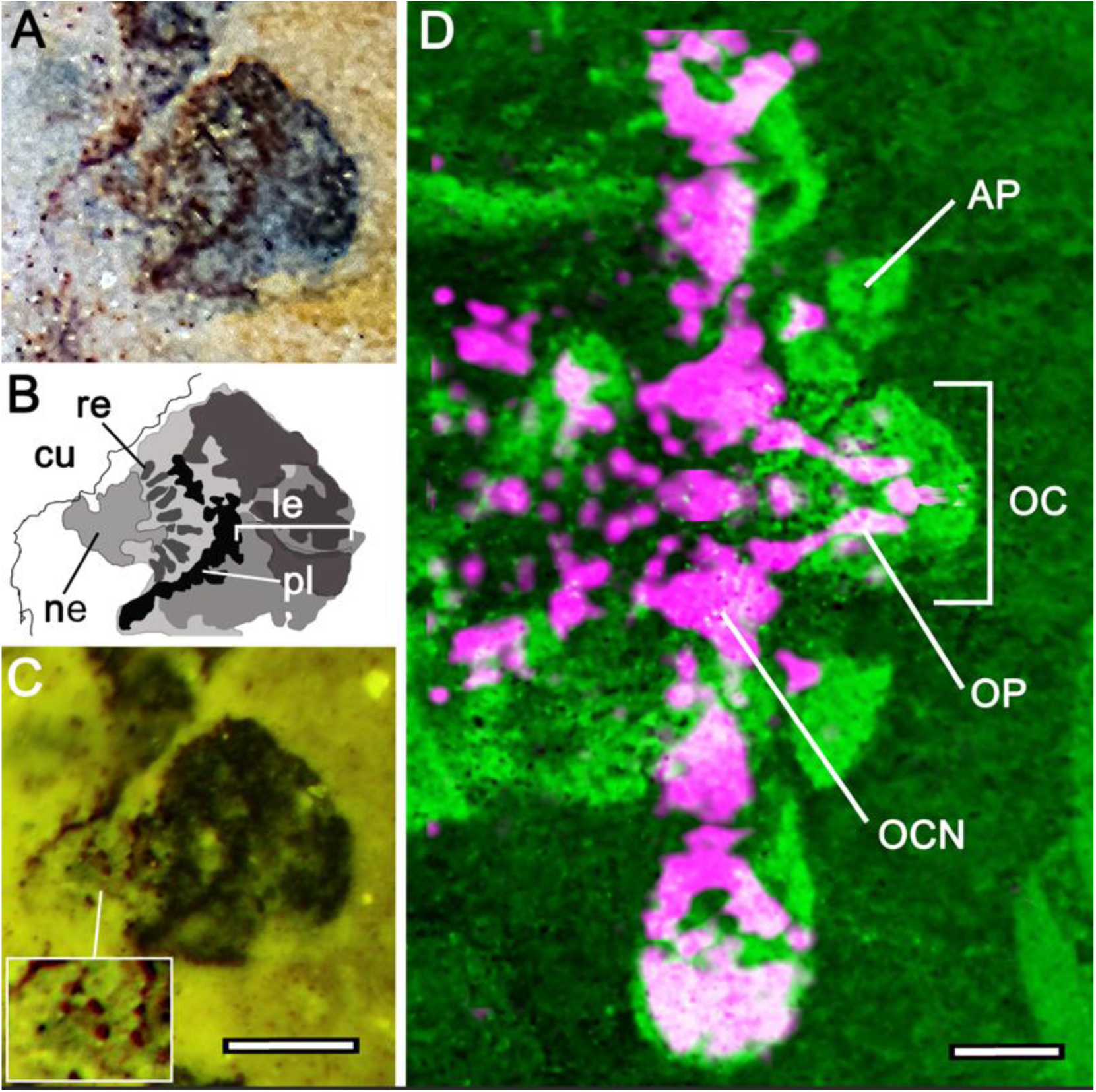
Concentric morphology of the anterior sclerite indicates a motile ocellar/nauplius- like “eye” in *Jianfengia*. Despite the compression of fossil specimen YKLP11117, its midline anterior sclerite (**A, B**) reveals concentric arrangements indicating outer dioptric elements interpreted as a triplet of lenses (le) overlying a pigmented layer (pl) beneath which are short columnar elements here interpreted as photoreceptors (re). The entire sclerite arises from a short neck (ne) (note panels A and B are identical to Fig. 1C, D). (**C**) UV illumination resolves the neck as comprising fold-like elements suggesting unsclerotized arthrodial membrane (inset lower left) extending from beneath the overlying frontal cuticle (cu). (**D**) Specimen YKLP17299 (magenta) superimposed on YKLP11117 (green) to demonstrate the match between neural features and the external morphology of *Jianfengia*. Ocellar neural pathways (OP) extending to the ocellar neuropils (OCN) confirm that the ocellus (OC) is a *bona fide* sensory organ. Traces of neuropil coinciding with the cuticular anterior projection (AP) suggest its possible sensory role. Scale bars are 0.25mm.

**Fig. S2.**
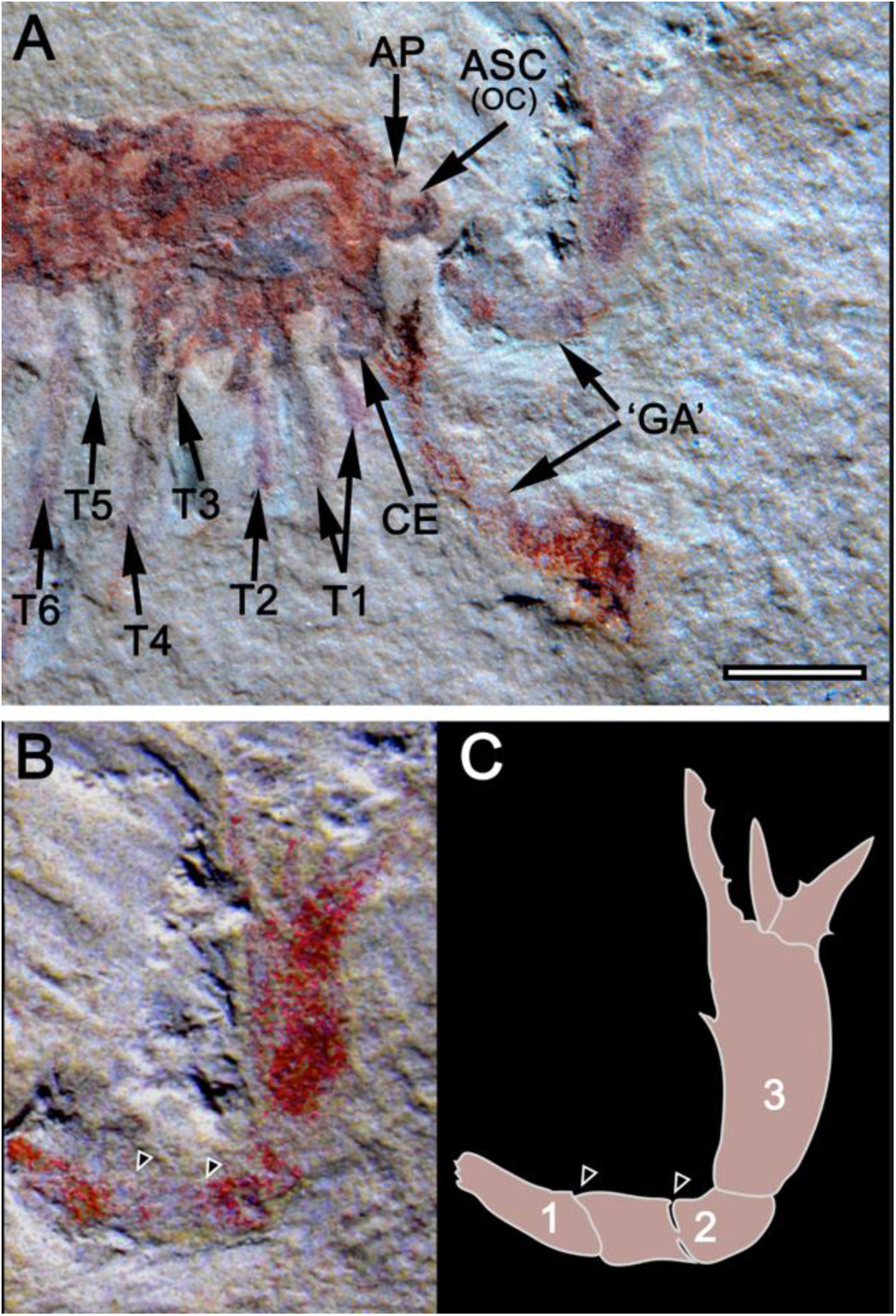
The great appendages of *Jianfengia*. (**A**) Lateral view of anterior appendicular arrangements of specimen NIGPAS 100123b. (‘GA’) Right and left ‘great appendages.’ The right compound eye (CE) is preserved, the left eye is buried in the matrix. The left anterior process (AP) is visible, its contralateral counterpart is obscured by matrix. The anterior sclerite carrying the ocelli [ASC (OC)] is visible. Both left and right biramous appendages of segment (T1) are in the same plane, whereas only the right appendages of segments T2-T4 are visible extending from beneath the carapace. (**B, C**) Isolated left GA showing podomeres (1, 2). Triangles indicate possible folds or sutures, which suggest flexibility within podomere 2 as well as between it and podomere 3. Scale bar for panels A-C=2mm.

**Fig. S3.**
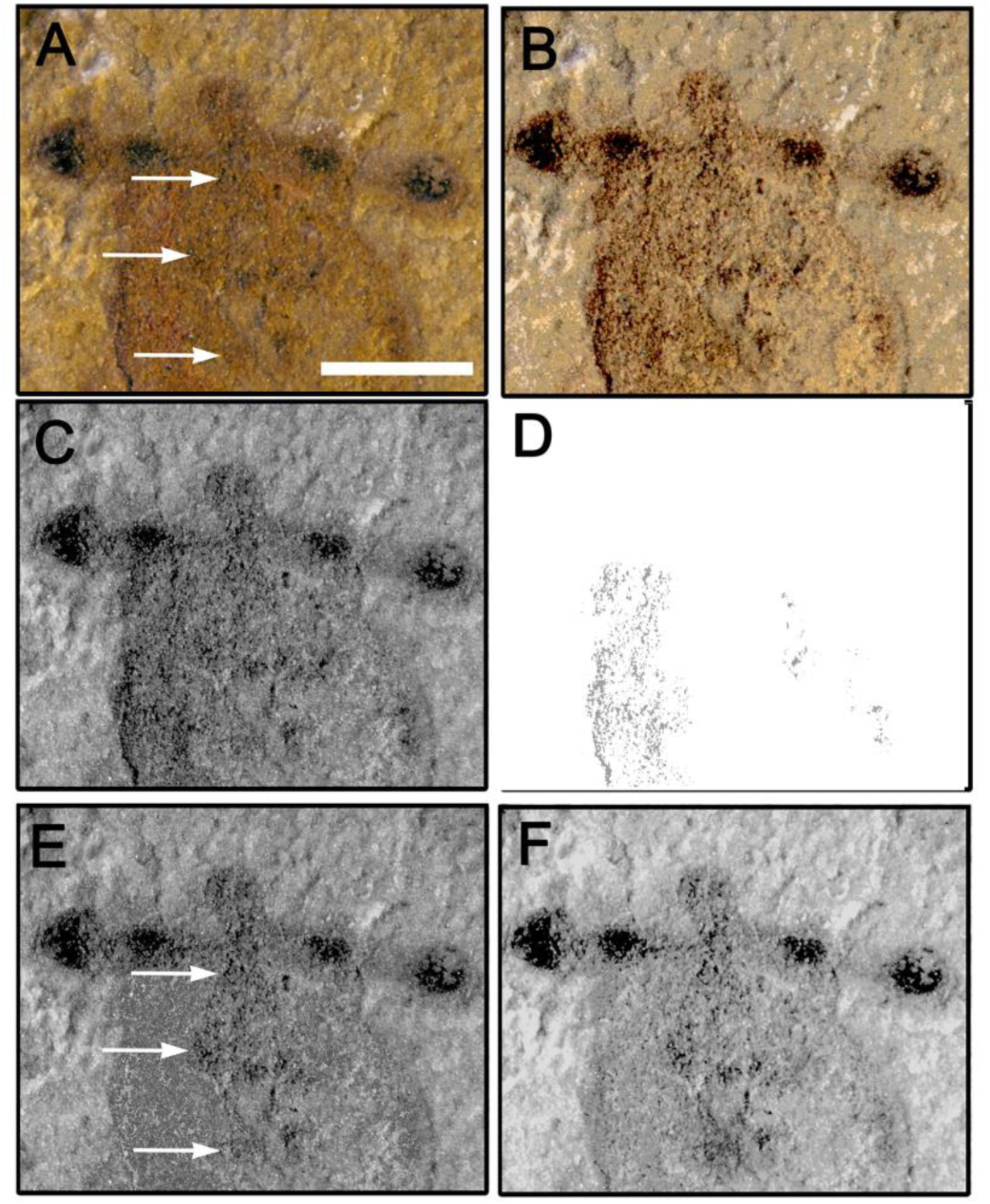
Exposure of trace neuropils and tracts in specimen YKLP 11367. (**A**) Photographic raw image reveals optic appendages and three mirror symmetric arrangements (arrowed in A) that can be made out despite color and density varying across the specimen. (**B**) Selective removal of the orange tint demarcates traces of fossilized soft tissue but also enhances the contrast of particulate surface features. (C-E). (**C**) These imbalances across the specimen can be rectified by converting the image to its greyscale mode. (**D, E**) Grey level densities responsible for imbalance (D) are merged with the relevant areas and thereafter adjusted to balance the left and right side of the fossil (E). (**F**) Adjustments of exposure and offset provide the final image for analysis and reconstruction (see Materials and Methods). Scale bar in panel A = 0.5mm

**Fig. S4.**
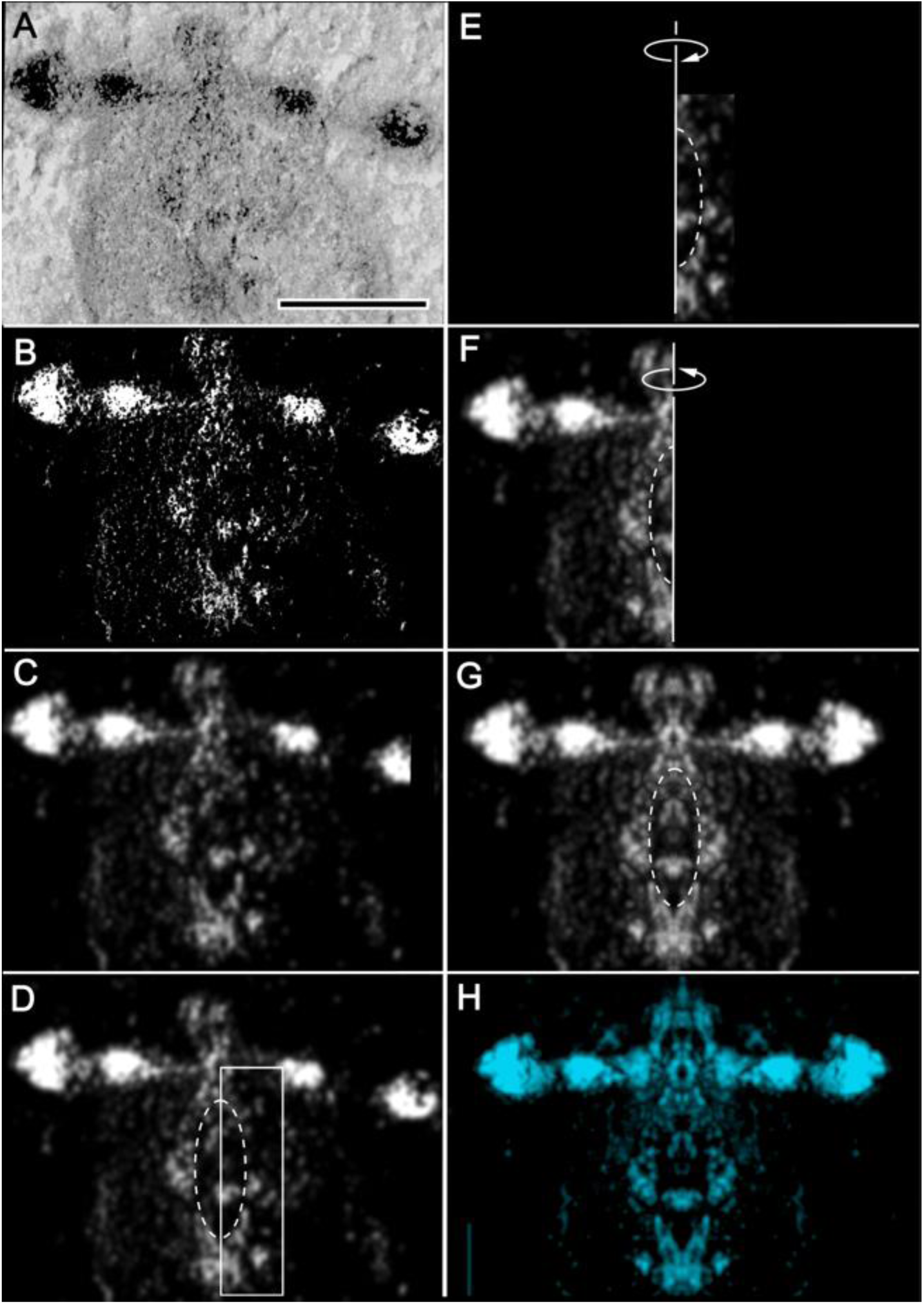
Reconstruction and interpretation of fossilized neural traces, neuropils, and connections of specimen YKLP11367. In panel A, the raw image shows a tilted specimen in which potential traces of fossilized tissue on the left are dissimilar to those on the right. This indicates that two levels of interest are exposed at or just beneath the surface of the fossil’s fracture plane. The material in panel A consists of distributed patches and granularities which clearly distinguish the more granular fossil against a paler background of the embedding matrix. Within the fossil, adjacent arrangements of dark particulate matter indicate residues of fossilized soft tissue. Following their isolation as described in panels B and C mirror symmetric overlay of both sides provide the summed depiction of a cerebral organization, the interpretation of which is enabled by reference to descriptions of eucrustacean brain neuroanatomy (see main text). (**A**) Raw gray level image; (**B**) Extracted image employing Adobe Photoshop functions to eliminate intensity levels below a defined threshold (87% black in standard CMYK scale). (**C**) Application of Gaussian blur function (R=10). (**D**) Selection of neural traces within the right half of the brain (boxed area) excluding the displaced right eye stalk and its contents. The stomodeum is indicated by the dashed ellipse. (**E**) Isolated area boxed in panel d. This is then flipped over to the other side so that it overlies the left half (**F**) using the stomodeal margins for alignment. The resultant composite left half is next duplicated and flipped back to the right side resulting in (**G**) a bilaterally symmetrical nervous system. (**H**) Final reconstruction with ocellar system of YKLP17299 superimposed for the interpretive Extended Data Fig. 3. Scale bar in A=0.5mm.

**Table S1.** Terminologies of euarthropod body partitioning and their underlying rationale. Various nomenclatures are used in naming parts of the brain, the segmented trunk, and their associated appendages with (*) segments T1-T4 ancestrally undifferentiated (33) but divergently specialized in many stem lineages, such as the acquisition of mandible morphology of the T2 basipod (30–32). Historical nomenclatures refer to traditional views of a segmented brain. However, its embryonic formation is asegmental and mediated by a developmental program that is genetically distinct from that specifying the reiterated segmentation of the trunk. The genetic code underlying nervous system partitioning (5, 45) provides the most parsimonious and biologically relevant terminology. Abbreviation: *s.l., sensu lato*.

| A-P axis partitioning | Segment term | Classic nomenclature | Genetic domain nomenclature | Associated appendage | Genetic code |
| --- | --- | --- | --- | --- | --- |
| 1<sup>st</sup> segment/<br>neuromere |  | Protocerebrum | ce1 domain =<br>prosocerebrum.<br>ce2 domain =<br>protocerebrum | ce1 = labra<br>ce2 =<br>eyestalks | • ce1=*FoxQ2-Six3-hbn*<br>• ce2=*Six3-Otx-Pax6*<br>• ce3=*Emx-Nk2-exd*<br>• Non-Hox<br>• Non-mesoderm |
| 2<sup>nd</sup> segment/<br>neuromere | A1 | Deutocerebrum | ce3 domain =<br>deutocerebrum | ce3 =<br>chelicerae<br>(*sensu.lato*),<br>antennules |  |
| 3<sup>rd</sup> segment/<br>neuromere | A2 | Tritocerebrum | Trunk segment T1 | biramous<br>limb/<br>appendage* | • Hox code<br>• mesoderm-related |
| Trunk<br>segment 1 | A3 | Mandibular<br>gnathal segment/<br>neuromere | Trunk segment T2 | biramous<br>limb/<br>appendage* |  |
| Trunk<br>segment 2 | A4 | 1<sup>st</sup> Maxillary<br>gnathal segment/<br>neuromere | Trunk segment T3 | biramous<br>limb/<br>appendage* |  |
| Trunk<br>segment 3 | A4 | 2<sup>nd</sup> Maxillary<br>gnathal segment/<br>neuromere | Trunk segment T4 | biramous<br>limb/<br>appendage* |  |
| Trunk<br>segment 4 | T1 | 1<sup>st</sup> trunk segment | Trunk segment T5 | biramous<br>limb/<br>appendage |  |
| Trunk<br>segment 5 | T2 | 2<sup>nd</sup> trunk segment | Trunk segment T6 | biramous<br>limb/<br>appendage |  |
| Trunk<br>segment 6 | T3 | 3<sup>rd</sup> trunk segment | Trunk segment T7 | biramous<br>limb/<br>appendage |  |
| Trunk<br>segment 7 | T4 | 4<sup>th</sup> trunk segment | Trunk segment T8 | biramous<br>limb/<br>appendage |  |
| ... | ... | ... | ... | ... |  |

**Table S2.**
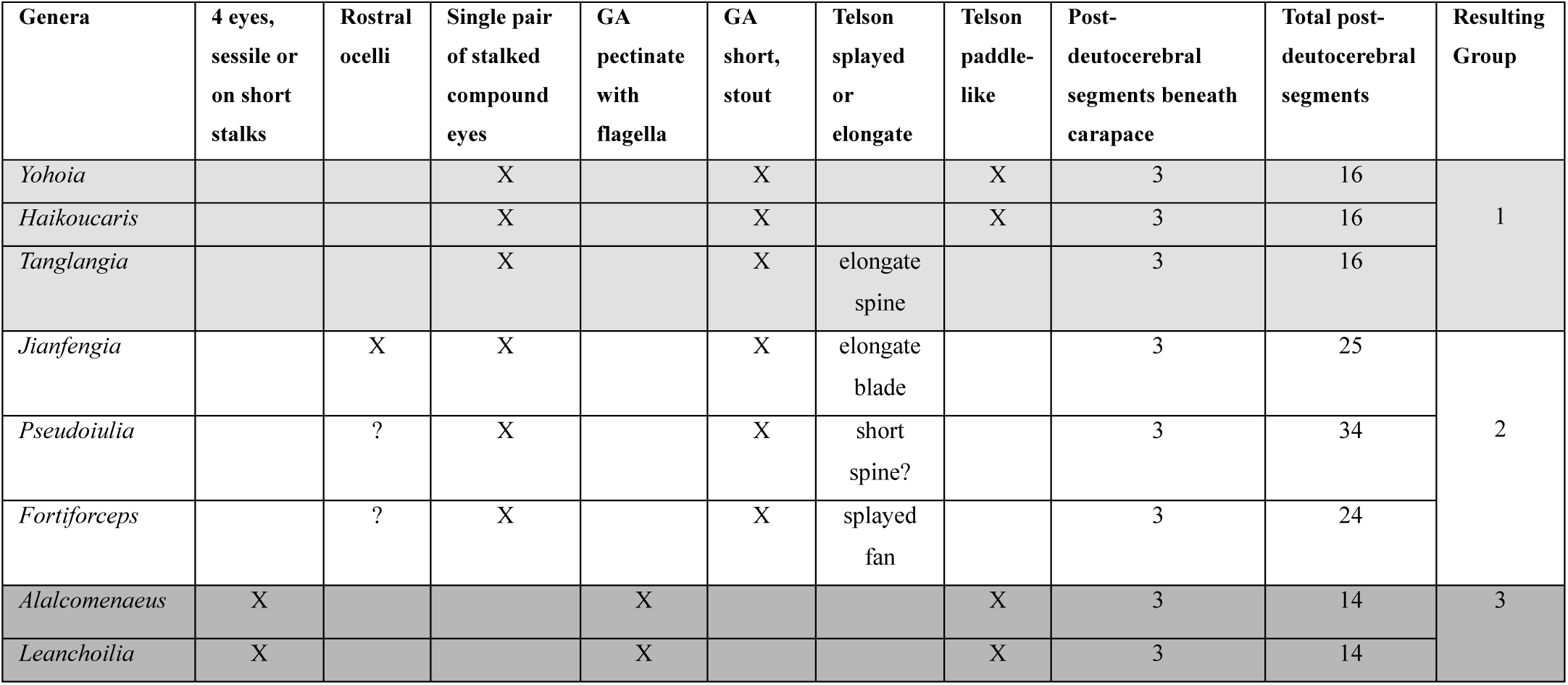
Subtype-specific features classify Megacheira. The paraphyletic euarthropod clade known as Megacheira is denoted by its deutocerebral appendages, also called “great appendages” (GAs) composed of two proximal articles with an elbow-like joint continuing as 3-4 distal articles, each providing an upward/medial spine. Genera listed here exclude novel specimens known from single examples. Cross-comparison of species-specific features identifies three natural groups based on combinations of three traits. GROUP 1 (*Yohoia, Haikoucaris, Tanglangia*): three post-deutocerebral trunk segments obscured by carapace, 16 trunk segments total; single pair of stalked compound eyes; robust GAs with stout spines. GROUP 2 (*Jianfengia, Pseudoiulia* (synonym *Sklerolibyon* (39)), *Fortiforceps*): three post-deutocerebral trunk segments wholly obscured by carapace, 24-34 trunk segments total; single pair of stalked compound eyes; spike-like distal articles of the stout GAs. GROUP 3 (*Alalacomeneaus, Leanchoilia*): three post-deutocerebral trunk segments obscured by carapace, 13 trunk segments total; quartet of sessile, or short-stalked eyes; delicate GA with three pectinate distal articles each providing a prolonged flagellum.

